# *Cryptosporidium* infection of human small intestinal epithelial cells induces type III interferon and impairs infectivity of Rotavirus

**DOI:** 10.1101/2023.08.30.555581

**Authors:** Valentin Greigert, Iti Saraav, Juhee Son, Denise Dayao, Avan Antia, Saul Tzipori, William H. Witola, Thaddeus S. Stappenbeck, Siyuan Ding, L. David Sibley

## Abstract

Cryptosporidiosis is a major cause of severe diarrheal disease in infants from resource poor settings. The majority of infections are caused by the human-specific pathogen *C. hominis* and absence of in vitro growth platforms has limited our understanding of host-pathogen interactions and development of effective treatments. To address this problem, we developed a stem cell-derived culture system for *C. hominis* using human enterocytes differentiated under air-liquid interface (ALI) conditions. Human ALI cultures supported robust growth and complete development of *C. hominis* in vitro including all life cycle stages. *C. hominis* infection induced a strong interferon response from enterocytes, likely driven by an endogenous dsRNA virus in the parasite. Prior infection with *Cryptosporidium* induced type III IFN secretion and consequently blunted infection with Rotavirus, including live attenuated vaccine strains. The development of hALI provides a platform for further studies on human-specific pathogens, including clinically important coinfections that may alter vaccine efficacy.

## INTRODUCTION

*Cryptosporidium* spp. are apicomplexan parasites that cause diarrheal disease in humans and animals. The primary species responsible for human infection are *C. parvum*, which can also infect agricultural ruminant animals and is zoonotic, and *C. hominis*, which is transmitted from person to person (1). Cryptosporidiosis is recognized as one of the leading causes of severe diarrhea in children under the age of two in developing countries, with lasting developmental defects observed even after infection subsides (2, 3). There is currently no effective vaccine for cryptosporidiosis, and the only FDA-approved drug for treatment, nitazoxanide, has limited efficacy in immunocompromised individuals and is not approved for use in young children (4). Therefore, a deeper understanding of parasite biology and host-pathogen interactions is essential for identifying potential therapeutic and vaccine targets.

Most infections in children in the developing world are caused by *C. hominis*, a human-specific pathogen that lacks robust systems for laboratory study. As a result, studies on *Cryptosporidium* have largely used *C. parvum* as a model due to its ability to be propagated easily using immunodeficient mice (5). Although some animal models, such as intratracheal rat or gnotobiotic piglet (6–8), are used to study *C. hominis*, they are costly and require veterinary expertise unavailable for many laboratories. Furthermore, despite a recent interest in more complex in vitro systems to study host-parasite interactions during cryptosporidiosis (9), most studies published over the past three decades have been based on colonic neoplastic cells, such as HCT-8, HT-29, or Caco-2 cells, which exhibit many distinct features from the host small intestinal epithelium including, but not limited to, a lack of diversity of cell types. Although several systems have been described for in vitro growth of *C. parvum* in human small intestine and lung organoids (10) or murine stem-cell derived intestinal epithelial cultures (11), neither of these systems has been shown to be capable of supporting growth of the human pathogen *C. hominis*.

Here, we adapted the system we had previously developed for propagating murine stem cells from the gut, and for differentiation into intestinal cell lineages in vitro, using a method referred to as Air Liquid Interface (ALI) (11). Human ALI (hALI) cultures fully differentiated into all the normal lineages and sustained robust *C. hominis* growth in vitro, including production of oocysts. *Cryptosporidium* infection of hALI induced a strong type III interferon (IFN) response and adversely impacted concurrent Rotavirus (RV) infection. The development of hALI provides a valuable tool to study the complex interaction between the intestinal epithelium and human specific pathogens, including co-infection studies.

## RESULTS

### The hALI system displays a high level of cell differentiation

We have previously described a murine air-liquid interface (ALI) system for differentiation of intestinal epithelial culture and long-term propagation of *C. parvum* (11). However, in initial testing, this system did not support the growth of the human-specific pathogen *C. hominis* (**Fig. S1**). We reasoned that this limitation might be due to host range restriction and therefore we sought to develop a similar system using human intestinal stem cells differentiated under ALI conditions (**Fig. 1A**). Stem cells originating from ileal crypts were first propagated as spheroids using protocols described previously (11).

**Fig. 1.**
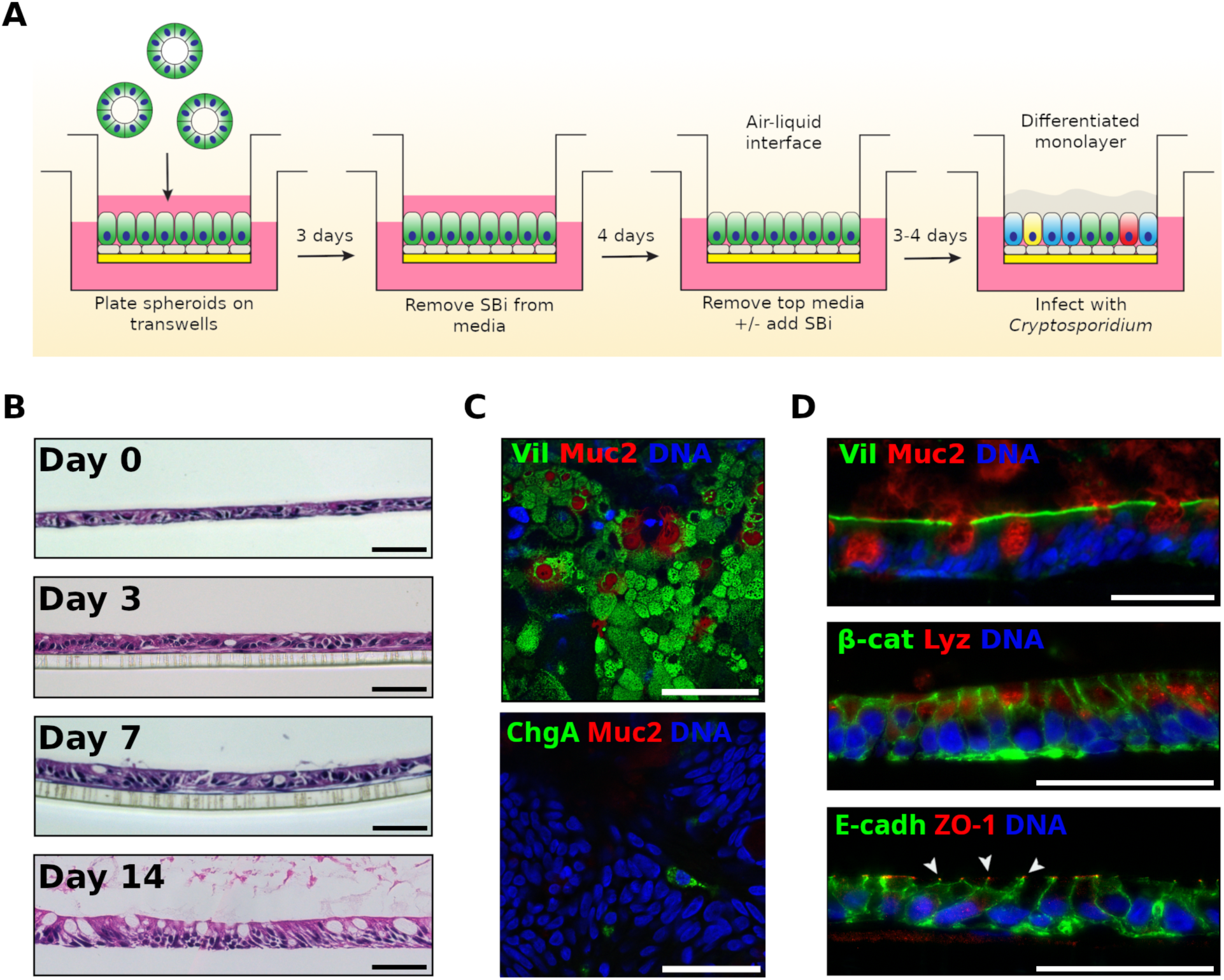
Human ALI system promoted development of a highly complex epithelial tissue composed of differentiated cell types. (A) Human air-liquid interface (hALI) was generated by seeding intestinal stem cells on a confluent layer of irradiated human intestinal myofibroblasts platted on Matrigel-coated transwells. Cells were cultured in 50% conditioned media (CM) complemented with Y-27632 (10 μM) and an inhibitor of the TGF-β pathway SB431542 (SBi, 10 μM). After 72 hr of growth, SB431542 was removed from the growth media. Seven days after seeding, medium from the top chamber was removed to generate ALI cultures, and SB431542 added back to culture medium when the purpose of the culture was parasite growth. Cells were used for infection 3 to 4 days after top medium removal. (B) Hematoxylin-eosin staining of sections of hALI showing the thickening and differentiation of the monolayer over time after removal of medium in the top chamber. Enterocytes became more columnar and goblet cells more abundant and mucus-producing over time. (C, D) Immunostaining of flat mounted (C) or transversal sections (D) 14 days after top medium removal revealed an uniform brush-border stained by anti-villin antibodies, disrupted by goblet cells secreting mucus stained with anti- Muc2 antibodies, and the presence of both enteroendocrine cells (anti-chromogranin A antibodies) and Paneth cells (anti-lysozyme antibodies). Β-catenin antibodies delineate cells, similar to E-cadherin (adehrens junctions). ZO-1 (tight junctions) is expressed at the apical end of cellular junctions, on the lumen side (arrowheads). Scale bars = 50 μm.

Spheroids were dispersed, plated on semi-permeable transwell inserts and allowed to develop under ALI conditions (**Fig. 1A**). ALI cultures were optimized using 50% conditioned medium complemented with rho-kinase inhibitor (Y-27632) and ALK4/5/7 inhibitor (SB431542) to promote growth and differentiation into monolayers resembling the physiological human ileal epithelium (**Fig. 1A**). After seven days of submerged culture, medium was removed from the top culture chamber triggering enhanced mucus production and differentiation. Characterization of ALI cultures by hematoxylin-eosin staining of transversal sections showed the cell layer became thicker after removing the medium from the top chamber as the cells appeared more columnar with time. Differentiation was apparent by an increased number of mucus-secreting goblet cells, particularly after 7 days of ALI culture (**Fig. 1B**).

Further characterization, using fluorescent markers for various intestinal cell types, revealed the existence of various cell types in the ALI culture. Enterocytes, displaying the brush border marked by villin staining, were the most common cell-type (**Fig. 1C, D**). Staining of the mucin-2 protein identified many mucus-secreting goblet cells (**Fig. 1C, D**). Other cell types also included Paneth cells, marked for lysozyme, and rare enterochromaffin cells marked by chromogranin A (**Fig. 1C, D**). The tissue showed the formation of adherens junctions (AJ), as shown by the expression and localization of E-cadherin at the interface between cells, and tight junctions (TJ), present between cells at the apical pole and marked by the presence of the ZO-1 protein (**Fig. 1D**).

Taken together, these results show that the human ALI (hALI) system achieves a high level of cellular differentiation, closely resembling in vivo human ileal epithelium.

### The hALI system supports extended *C. hominis* growth and oocysts formation

Having established a platform for hALI culture that mimics the small intestine, we used the system to culture *C. hominis*, a human-specific parasite. We modified the conditions slightly to enhance cell proliferation by treatment with the TGF-β pathway inhibitor SB431542 (see methods). Under these conditions, hALI supported substantial growth of *C. hominis* and *C. parvum* for up to 9 days (**Fig. 2A**). During this time period, parasite genome numbers increased from 100-300 fold. To explore the development of *C. hominis* in culture, we used a panel of antibodies developed against intracellular stages of *C. parvum* (12). Preliminary analysis verified that these regents cross-react with *C. hominis* stages (**Fig. S2**). By selecting a panel of mAbs that recognize stages across the life cycle, we showed that *C. hominis* underwent all stages of its life cycle during ALI culture, including the formation of gamonts (**Fig. 2B**). We were also able to detect newly formed oocysts recognized by surface staining with a commercial antibody called Crypto-a-Glo (**Fig 2 C, D**). To distinguish newly formed oocysts from the input inoculum we used EdU staining, which only stains proliferating nuclei (**Fig. 2C, D**). Oocysts that were positive for EdU staining contained four newly formed nuclei and a surrounding oocyst wall indicating that both *C. hominis* and *C. parvum* were capable of completing the entire life cycle in the hALI system. Hence, hALI, provides a valuable system to study human-specific pathogens, including anthroponotic *Cryptosporidium* strains and species.

**Fig. 2.**
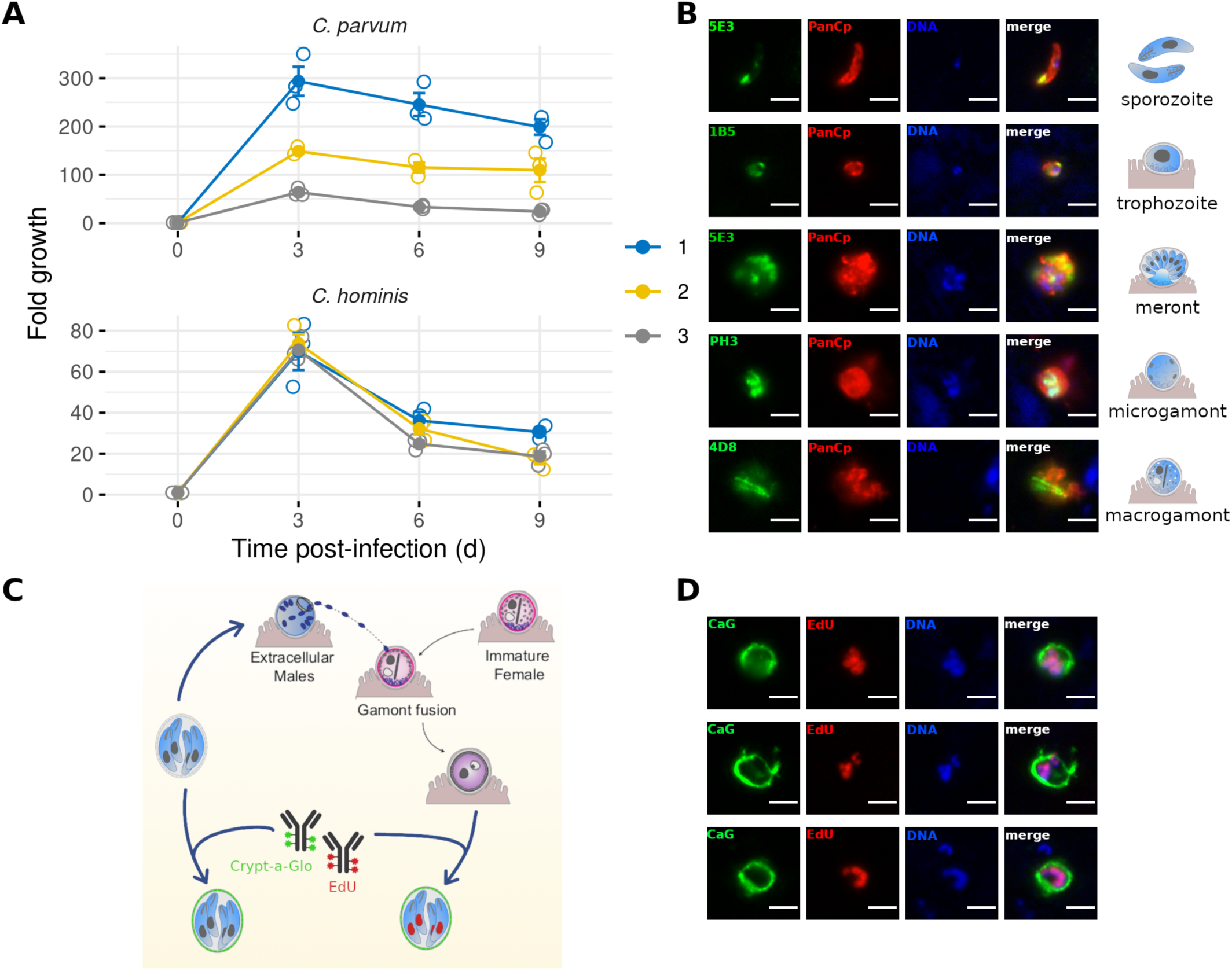
Human ALI system supported *Cryptosporidium hominis* culture with complete life cycle development. (A) Growth of *C. hominis* in hALI cultures over a 7-day period. Mean ± S.E. N = 3, n = 3 (B) Immunostaining of various stages of *C. hominis* cultured on hALI using a panel of previously described mAb to *C. parvum* (green) and a pan-*Cryptosporidium* polyclonal antibody (red)(12). Sporozoites and trophozoites were stained 6 hr after infection (pi), meronts 24 hr pi and microgamonts and macrogamonts 72 hr pi. (C) EdU was used in combination with Crypt-a-Glo (CaG) to differentiate newly-formed oocysts from carryover oocysts. (D) Immunostaining of newly-formed oocysts following infection of hALI with *C. hominis* oocysts, 72 hr pi. Scale bar = 3 μm

### Infection of hALI with *Cryptosporidium* species does not disrupt barrier function

Previous studies using adenocarcinoma cells have suggested that infection with *Cryptosporidium* induces loss of barrier function (13–16). Since hALI provides a monolayer composition that more closely mimics the intestine in vivo, we chose to re-examine the effect of *Cryptosporidium* infection on barrier function. We observed that 24 hr infection with either *C. parvum* or *C. hominis* did not alter the barrier function of hALI using both Trans-Epithelial Electrical Resistance (TEER) measurements and dextran flux assays (**Fig. 3A**). Similar experiments using HCT-8 cells resulted in strong alterations of barrier function following infection with *Cryptosporidium* (**Fig. 3A**), consistent with previous studies (14, 15).

**Fig. 3.**
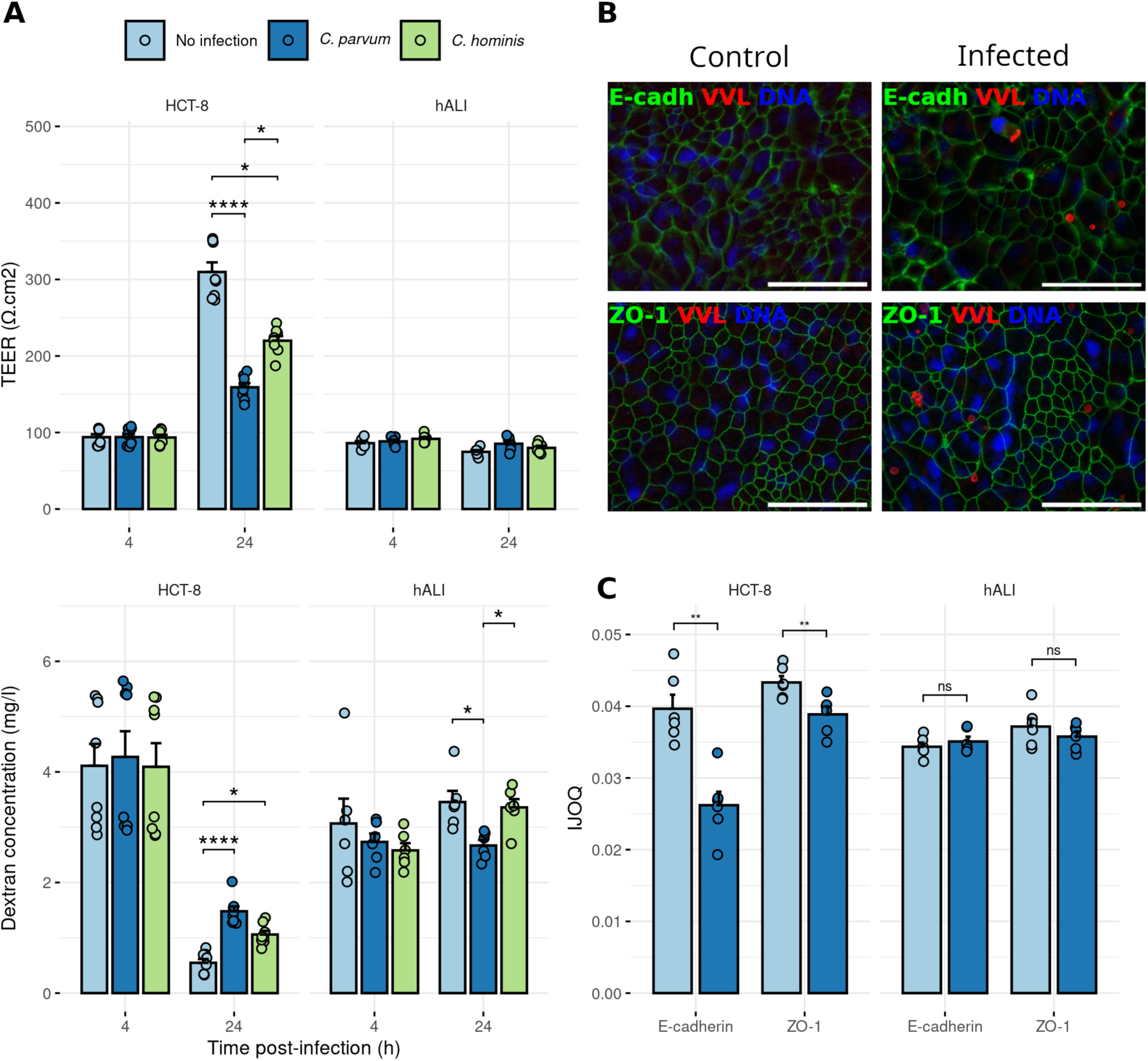
The permeability of human ALI did not increase after infection, as opposed to HCT-8 cell monolayers. (A) Trans-epithelial electrical resistance (TEER) and Dextran flux measured across hALI of HCT-8 monolayers infected with *C. parvum* or *C. hominis.* hALI was infected 3 days after top medium removal. Mean ± S.E. N = 2, n = 4. Kruskal-Wallis test followed by Dunn’s multiple comparison test. * *P* < 0.05, **** *P* < 0.0001. (B) Immunostaining of flat mounted hALI transwell culture showed tight junction (TJ; ZO-1, green) and adherens junction (AJ; E-cadherin, green) organization, with or without infection with *C. parvum* (*Vicia villosa* lectin, VVL, red). Scale bar = 50 μm. (C) Intercellular junction organization quantification (IJOQ) of hALI and HCT-8 cells infected with *C. parvum vs.* not infected, for AJ (E-cadherin) and TJ (ZO-1). hALI was infected 3 days after top medium removal. One representative experiment for each cell type is showed. Mean ± S.E. N = 2, n = 6. Mann-Whitney U test. ** *P* < 0.01, ns = non significant.

To further confirm the differences seen between hALI and HCT-8 cells, we examined the organization of both TJ and AJ, by staining the proteins ZO-1 and E-cadherin, respectively. Confocal imaging of these samples showed no disruption of TJ or AJ in hALI monolayers following infection with *Cryptosporidium,* as staining for the proteins consistently delineated the border of the cells in both conditions (**Fig. 3B**). In contrast, infection of HCT-8 cells resulted in lower expression of the ZO-1 and E-cadherin overall, and a loss of border definition of ZO-1 or E-cadherin staining (**Fig. S3**).

Quantification of the organization of TJ and AJ using the Intercellular Junction Organization Quantification (IJOQ) software, showed no alteration of junctional organization following infection in hALI, contrary to what was observed in HCT-8 cells (**Fig. 3C**). Collectively, these results demonstrate that infection of hALI cultures with *Cryptosporidium* does not displace TJ and AJ proteins and does not alter barrier function.

### Infection of hALI with *Cryptosporidium* upregulates IFN pathways

To further explore the hALI response to *Cryptosporidium*, we designed an RNA-Sequencing (RNASeq) experiment to investigate pathways that are perturbed by infection. hALI cultures were infected with either *C. hominis, C. parvum*, or as a mock-infected control and allowed to grow for up to 72 hr (**Fig. 4A**). Cells were harvested 24 hr after infection for bulk RNA-Seq. In parallel, cultures medium was collected from the bottom chamber for determination of soluble secreted chemokines at 4 hr, 24 hr, and 72 hr after infection (**Fig. 4A**).

**Fig. 4.**
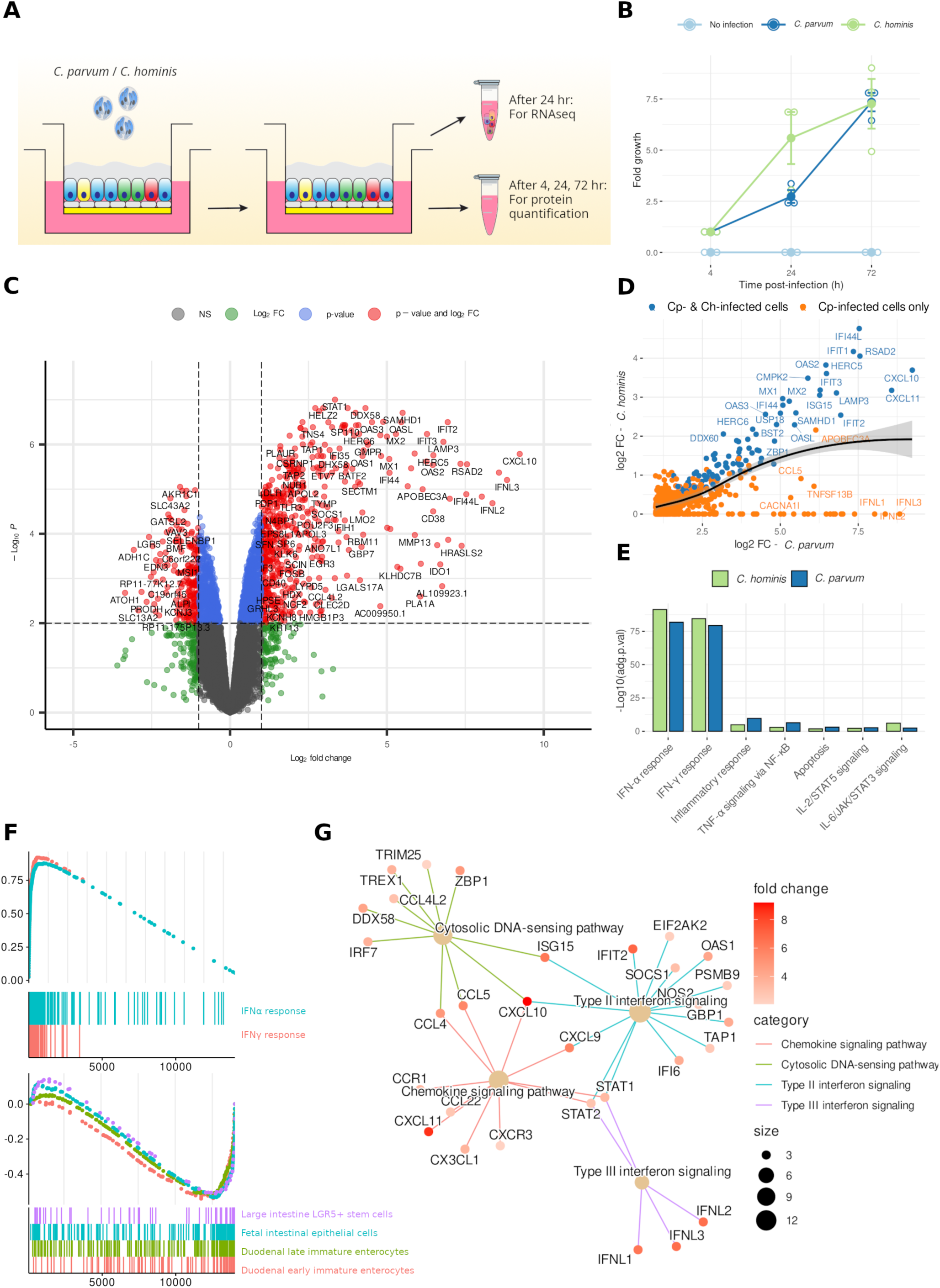
Infection of human ALI with *Cryptosporidium* induced the expression of genes involved in inflammation, including type I, II and III interferon pathways. (A) hALI was infected with either *C. parvum* or *C. hominis*, with a mock infection as control, for a total of 72 hr. Cells were harvested for RNA-Seq after 24 hr of infection. Bottom chamber media was harvested for cytokine quantification using a Luminex bead assay after 4 hr, 24 hr and 72 hr of infection. (B) Growth of *C. hominis* and *C. parvum* in hALI cultures used for this experiment over a 3-day period. Mean ± S.E. N = 3, n = 1. (C) Volcano plot of differentially expressed genes in hALI 24 hr after infection with *C. parvum*. Genes up- or down-regulated by more than 2-fold and *P* < 0.05 are shown in red. (D) Scatter-plot showing genes upregulated following *C. hominis* infection vs. *C. parvum* infection. (E) Pathway analysis of upregulated genes following *C. hominis* or *C. parvum* infection (MsigDB Hallmark 2020). (F) GSEA analysis showing upregulated (up) and down-regulated (down) pathways in hALI infected with *C. parvum*. (G) Network plot showing the involvement of highly expressed genes following *C. parvum* infection in several upregulated pathways (WikiPathway).

We observed similar parasite growth for both *C. parvum* and *C. hominis* over the course of the experiment, with a slight advantage for *C. hominis* after 24 hr of infection (**Fig. 4B**). RNA-Seq analysis showed the upregulation of many genes involved in inflammatory processes following the infection with either *C. parvum* or *C. hominis* (**Fig. 4C & Fig. S4**). Infection with either species caused a similar qualitative host response, primarily inducing type I and II IFN pathways, although the magnitude of induction and number of upregulated genes was higher following infection by *C. parvum* (**Fig. 4D-G & Table S1**). Additionally, infection with *C. parvum* resulted in downregulation of a number of genes, that were not observed following infection with *C. hominis*. These genes were found to be mainly expressed by immature intestinal epithelial stem cells, and included those involved in intestinal differentiation pathways, such as *ATOH1* or *BMP3* (**Fig. 4C-F**).

To further validate these results, we examined protein secreted into the culture medium of ALI using a Luminex bead-based immunoassay. The secretion of CXCL10 was enhanced following infection by either *C. parvum* or *C. hominis*, whereas secretion of CCL5 was only increased when *C. parvum* was used (**Fig. 5**). Although there was no significant upregulation of the type I IFNs when tested at the protein level, IFN-λ1 and IFN-λ2/3 were upregulated by *C. parvum* infection (**Fig. 5**).

**Fig. 5.**
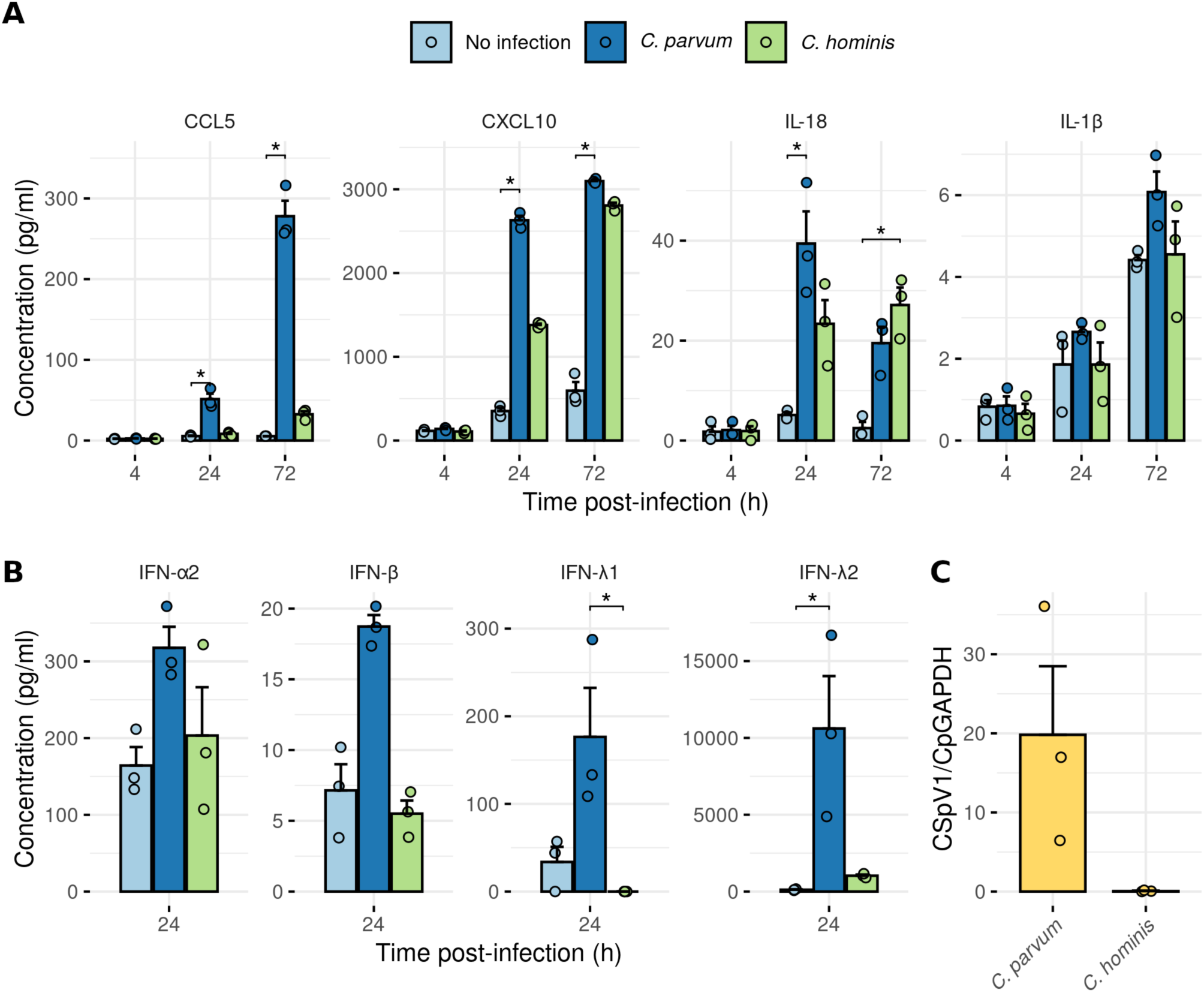
Infection of human ALI with *C. parvum* induced the production of inflammatory cytokines, including type I and III interferons. (A, B) Induction of chemokines and cytokines following infection with *C. parvum* or *C. hominis* based on Luminex bead assay. Mean ± S.E. N = 3, n = 1. Kruskal-Wallis test followed by Dunn’s multiple comparison test. * *P* < 0.05. (C) CSpV1 load in each batch of *Cryptosporidium* oocysts was measured by RT-qPCR. Each strain was tested thrice, on 3 different batches (i.e. 3 different oocyst-producing calves for *C. parvum*, and 3 different oocyst-producing gnotobiotic piglets for *C. hominis*).

These findings parallel the stronger induction of immune pathway genes by *C. parvum* infection as seen in RNA-Seq studies. Finally, we tested the level of secretion of interleukins IL-1β and IL-18, which are involved in the NLRP6 inflammasome, recently described as a key mechanism for enterocyte’s defense against *Cryptosporidium* infection in a mouse model (17). However, in our system, there was no increase in IL-1β secretion, although there was a strong increase in IL-18 secretion (**Fig. 5B**).

Taken together, these results show that *C. parvum* and *C. hominis* induce similar inflammation and IFN pathways in hALI cultures, although the magnitude of induction is much stronger in *C. parvum* infected cultures. Additionally, it is notable that *C. parvum* infection down-regulated differentiation of secretory lineages, consistent with previous findings that partially differentiated epithelium seems to be optimal for in vitro growth (11).

### *Cryptosporidium*-induced inflammation correlates with CSpV1 load in the parasite

*Cryptosporidium* contains a double-stranded RNA (dsRNA) virus called *Cryptosporidium parvum virus 1* (CSpV1) (18, 19). We hypothesized that CSpV1 might be involved in the induction of the observed responses, and that differences in the viral load between the two cryptosporidial species might explain the difference in amplitude of inflammatory response observed. Using RT-qPCR, we showed that the *C. parvum* strain used for our experiments contained substantial levels of CSpV1, with some batch-to-batch variability, whereas CSpV1 was consistently low or absent in *C. hominis* (**Fig. 5C**). These correlative findings suggest CSpV1 may be involved in the induction of inflammation observed following infection with *Cryptosporidium*.

### IFN induction by hALI infection with *C. parvum* inhibits RV infection

During early development, infants are often infected with multiple diarrheal agents, including bacteria, protozoa, and viruses (2). The strong induction of a type I/III IFN response following infection with *C. parvum* led us to consider what the physiological role this response might have on co-infection with enteric viruses. For this, we chose to look at the infection with RV, a pathogen epidemiologically related to *Cryptosporidium* (2) and known to be sensitive to the antiviral activities of type III IFN(20). For these experiments, hALI was infected with *C. parvum* for 24 hr, then infected with RV for 24 hr (**Fig. 6A**). Cells were harvested 2 hr and 24 hr after RV infection for viral genome copies and human IFNL3 mRNA quantified. As expected, IFNL3 expression was strongly upregulated in hALI infected with *C. parvum*, as opposed to non-infected hALI (**Fig. 6B**). RV infection alone also induced high levels of IFNL3 expression as expected (**Fig. 6B**).

**Fig. 6.**
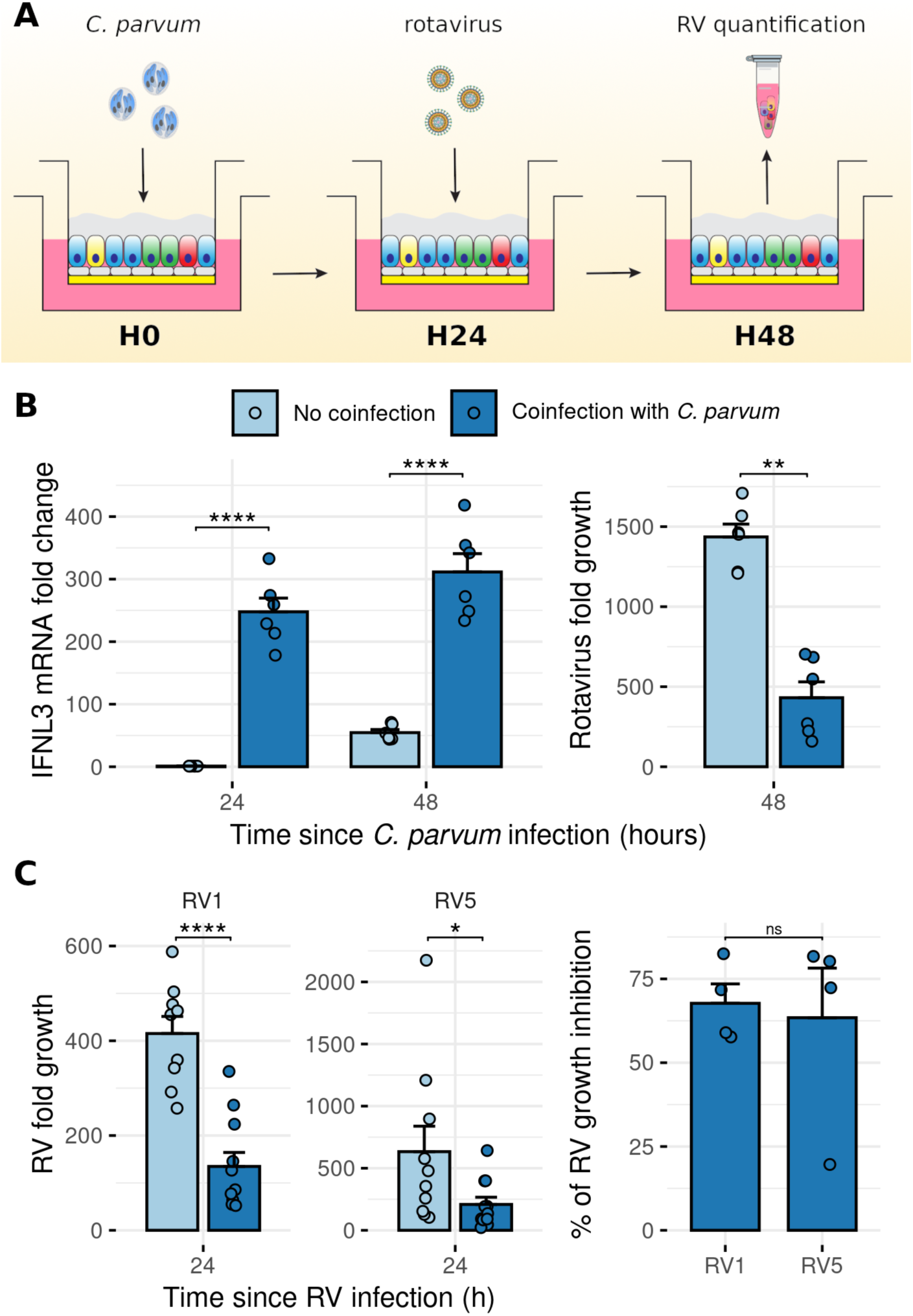
Pre-infection of human ALI with *C. parvum* inhibited Rotavirus growth. (A) hALI was infected with *C. parvum* for 24 hr prior infection with Rotavirus (RV). Cells were harvested 2 hr and 24 hr after infection with RV for virus quantification by RT-qPCR. H0, H24, H48: time since *C. parvum* infection (hr) (B) (left) IFN-λ3 gene expression as measured by RT-qPCR after hALI infection with *C. parvum* + RV *vs.* mock + RV infection. Mean ± S.E. N = 2, n = 3. One-way ANOVA followed with Tukey’s HSD test. (right) RV growth in hALI pre-infected with *C. parvum* as compared to uninfected hALI. N = 2, n = 3. Mann-Whitney U test. ** *P* < 0.01. (C) (left) RV vaccine strains growth inhibition in hALI pre-infected with *C. parvum* as compared to uninfected hALI. Mean ± S.E. N = 4, n = 3. After detection and removal of outliers with the 1.5xIQR rule, Mann-Whitney U test. * *P* < 0.05, **** *P* < 0.0001. (right) Percentage of inhibition of RV vaccine strains growth over 4 independent experiments. Mean ± S.E. Mann-Whitney U test. ns = non significant.

Importantly, quantification of RV copy number showed that viral growth was significantly inhibited in hALI that had been previously infected with *Cryptosporidium* (**Fig. 6B**). Because currently available RV vaccines are based on live attenuation strains that are avirulent due to genetic reassortment, we conducted similar experiments using both Rotarix (RV1) and RotaTeq (RV5), from GSK and Merck, respectively. Consistently, hALI infected with *C. parvum* 24 hr prior to RV infection led to a 60-70% inhibition of virus growth, an effect that was similar for both vaccine strains (Fig. 6C). This result shows that the host responses modulated by human intestinal epithelium infection with *Cryptosporidium* are substantial enough to impact the course of another infection caused by a completely different pathogen, including RV vaccine strains.

## DISCUSSION

Cryptosporidiosis is a major enteric disease in young children, the primary disease burden being due to *Cryptosporidium hominis*, which lacks tractable in vitro systems to for study. To address this deficiency, we developed a human stem cell derived culture system that sustained *C. hominis* growth through its entire life cycle in vitro. In hALI cultures, human epithelial cultures underwent differentiation of all lineages normally present in the small intestine and supported all stages of *C. hominis* growth, including the formation of new oocysts. Following infection with *C. hominis* or *C. parvum*, barrier function was not impacted, despite a strong inflammatory response primarily based on the activation of type I/III and II IFN pathways. This strong host response to *Cryptosporidium* infection inhibited the growth of RV, including several vaccine strains. The hALI system offers a platform to study human-restricted pathogens, to explore host pathogen interactions and to examine co-infection with human-specific pathogens.

We based our hALI system on previous work from our laboratory, in which we were able to propagate *C. parvum* in vitro in murine ALI cultures over a period of several weeks (11). Culture of stem-cell derived enterocytes under ALI conditions has previously been shown to be critical for cell differentiation and the ability of the tissue to sustain continued *C. parvum* growth (11). However, the previous system based on mouse intestinal stem cells does not sustain *C. hominis* growth in vitro. This mirrors what is seen in animal models where *C. parvum* has a broad host range and will infect ruminants to mice, while *C. hominis* is unable to do so (1). Whether restricted host range results from a difference in initial infection (i.e. a different in receptor or entry pathway) or the ability to productively expand is currently unknown. To accommodate the narrow host range of *C. hominis*, we expanded human intestinal stem cells as spheroids and differentiated them under ALI conditions. Human ALI cultures developed enterocytes, Paneth cells, goblet cells, enterochromaffin cells, strongly produced mucus, and developed well-defined architecture with a clearly stained brush border. The hALI system was capable of achieving this state without using differentiation-enhancing media, as has been described previously in other systems (21). By further modifying the hALI system to enhance cell proliferation by addition of a TGF-β pathway inhibitor, we were able to promote the growth of *C. hominis*, which underwent all stages of the life cycle, including the generation of new oocysts. Although previous organoid derived systems have also been described for in vitro culture of *C. parvum* (11, 22, 23), none of these have shown utility for growth of *C. hominis*. The development of hALI provides a platform that might also be useful for other human-specific pathogens.

Previous in vitro and in vivo studies have shown that infection with *Cryptosporidium* results in an increased permeability of the epithelium (13–16, 23, 24). Several mechanisms have been proposed to explain this consequence of infection, including the release of IL-1β and TNFα by monocytes (14), the alteration of expression of TJ and AJ proteins (15), or the induction of autophagy in intestinal epithelial cells (IEC) (16). We also observed disruption of junctions in HCT-8 cells, although despite robust infection, we were surprised to observe that hALI did not exhibit altered barrier function or distribution of AJ and TJ proteins. Adenocarcinoma cell lines, such as Caco-2 and HCT-8 cells, that are widely used to study cryptosporidiosis, also display decreased specific cell-cell adhesion, which likely facilitates metastasis to reach other organs (25). Hence, these adenocarcinoma models likely do not represent the response of normal healthy tissue. In contrast to our findings, infection of stem-cell derived enterocytes cultured with continuous medium coverage (i.e. submerged), did display barrier function alterations following infection with *Cryptosporidium*, with the disorganization of both TJ and AJ (23). This difference may result from physical differences between submerged cells bearing the weight of media as opposed to hALI being protected by a layer of mucus, or other differences in the protocols such as the burden of infectious challenge. Other studies have reported barrier disruption in vivo due to the release of IL-1β and TNFα by monocytes (14). The hALI model would not be expected to reproduce this feature as monocytes are missing from monolayers. Finally, in vivo studies have been conducted with neonatal mice where compromised barrier function has been seen (24). Whether barrier disruption is also a predominant feature of infection in young children is uncertain, but it may play an important role in the disease severity. Our in vitro studies suggest that barrier disruption is not a direct consequence of infection but more likely ensues from the inflammatory immune response that occurs in vivo (26).

Infection with either *C. parvum* or *C. hominis* induced strong activation of inflammatory processes, primarily due to the expression of genes belonging to both type I/III and type II IFNs. Type II IFN, specifically IFN-γ, has been shown to play a protective role in the innate immune response to *Cryptosporidium* infection in rodents (5, 6, 27–31), in vitro cell lines (32, 33), and humans (34–36). Recent research using a mouse model of cryptosporidiosis demonstrated that the production of IFN-γ by innate lymphoid cells (ILC) was essential for controlling the parasite and was stimulated by the synergistic effect of interleukin (IL)-12 and IL-18 secreted by epithelial cells (31), as suggested by previous studies (37). We observed enhanced secretion of IL-18 following *Cryptosporidium* infection, without upregulation of the IL-18 gene, suggesting the cleavage and release of preformed pro-IL-18 (38). It is likely that enterocyte-derived IL-18 is released due to NLRP6 inflammasome activation by *Cryptosporidium*, a mechanism that was recently described (17). Our findings are also consistent with a previous study using human small intestine and lung organoids that demonstrated that infection with *C. parvum* upregulated genes associated with type I IFN immunity (22). Type I IFNs have been shown to be involved in the innate immune response to *Cryptosporidium* infection with a protective effect (39). Type I and type III IFN pathways are highly overlapping, and the later pathway is likely more important in intestinal epithelium, which primarily expresses the receptors for IFN-λ (40). Consistent with this idea, two recent studies have shown a role for type III IFNs during cryptosporidiosis, mitigating the alteration of barrier function in various in vitro and in vivo models (24) and acting as a protective factor against *Cryptosporidium* infection in a STAT1-dependent manner, mediated by the pattern recognition receptor TLR3 (41).

One of the major differences observed between *C. parvum* and *C. hominis* infections in hALI was the absence of upregulation of the *IFNL1*, *IFNL2*, and *IFNL3* genes, or downstream proteins, in the latter. This difference might reflect the fact that while the pathways induced by *C. parvum* and *C hominis* were similar, the magnitude of responses was quite different. *Cryptosporidium* hosts a dsRNA virus called *Cryptosporidium parvum virus 1* (CSpV1) (18, 19). Hence, a difference in CSpV1 load between the two species might explain the observed difference in induction of type III IFN, as well as the difference in CCL5 expression, which is regulated by the IRF3 pathway downstream of TLR3 (42). Furthermore, CSpV1 has recently been involved in cryptosporidiosis pathophysiology by inducing the type I IFN pathway, inhibiting in turn the IFN-γ pathway, providing a form of protection against the immune response to the parasite (43). It is worth noting that in this study, type I IFN activation by CSpV1 was not dependent of TLR3, but instead on the Pkr and Rig-I/Mavs/Sting signaling pathways (43). In any case, further exploration of the role of CSpV1 in cryptosporidiosis pathophysiology, using more direct methods, is warranted.

Following *C. parvum* infection, several genes involved into intestinal cells differentiation were downregulated, such as *ATOH1* and *BMP3*. A previous study showed that these secretory lineages are not necessary for propagation of *C. parvum* in murine ALI, and that partially differentiated epithelium seems to be optimal for in vitro growth of the parasite (11). Together, these findings suggest that downregulation of differentiation enhancing genes might be part of a parasite strategy to limit its host cell differentiation to benefit longer from a more suitable niche. Host gene manipulation by various apicomplexan parasites has been frequently described (44). However, intestinal inflammation can also impair differentiation (45, 46). Thus, whether the downregulation of differentiation genes is a consequence of the inflammation or of a direct manipulation by the parasite remains to be elucidated.

Rotavirus, a type of non-enveloped dsRNA viruses from the Reoviridae family, is a major cause of dehydrating diarrhea in young children, particularly in intertropical areas, similar to cryptosporidiosis (2, 47). Extensive research efforts have led to the development of RV vaccines and globally widespread vaccination campaigns (47). However, while the efficacy of these vaccines is high in wealthy countries such as the USA, it is lower in countries with high child mortality rates, i.e. in lower income settings, correlating with a lack of universal access to clean water (47–49). We demonstrated that infection of hALI with *Cryptosporidium* significantly reduced RV growth, including the vaccine strains RV1 and RV5. *Cryptosporidium* infection led to an increased production of IFN-λ3, ultimately resulting in the inhibition of RV growth. These findings raise the intriguing hypothesis that co-infections, such as cryptosporidiosis, may modify the inflammatory state of the intestinal epithelium and adversely impact the efficacy of vaccines (48). Rotavirus and cryptosporidiosis co-exist in the same localities, hence such co-infections are likely to occur in young infants in resource poor settings and this may partially explain the variable efficacy of vaccines from different regions.

Cryptosporidiosis remains a primary health concern in the developing world, with many unanswered questions relating to both parasite’s biology and host response to the infection. Tools allowing to study these aspects of human cryptosporidiosis are therefore needed to better explore host-pathogen interactions using systems closely mimicking the physiological human intestinal epithelium. The hALI system thus provides a valuable platform to deepen our understanding of the parasite, the disease, co-infections, and intestinal homeostasis.

## EXPERIMENTAL MODEL AND SUBJECT DETAILS

### Cryptosporidium strains

*C. parvum* oocysts (AUCP-1 isolate) were maintained by repeated passage in male Holstein calves and purified from fecal material after sieve filtration, Sheather’s sugar flotation, and discontinuous sucrose density gradient centrifugation as previously described (50). All calf procedures were approved by the Institutional Animal Care and Use Committee (IACUC) at the University of Illinois Urbana-Champaign. Purified oocysts were stored at 4°C in PBS + 50 mM Tris-10 mM EDTA, pH 7.2 for up to six months before use.

The *C. hominis* strain TU502 was originally isolated from a child in Uganda (7, 51). TU502 isolate was maintained and serially passaged in gnotobiotic piglets at Tufts University. Oocysts were purified from feces collected daily from infected piglets on Nycodenz step gradients (Alere Technologies AS) as described previously (52). All piglet procedures were conducted in compliance with the study protocol G2020-97, approved by the Institutional Animal Care and Use Committee at Tufts University Cummings School of Veterinary Medicine in accordance with the Guide for the Care and Use of Laboratory Animals (National Research Council). Purified oocysts were stored at 4°C in sterile PBS until use.

### Cell lines

All cell lines were cultured at 37°C in a 5% CO_2_ incubator under normal atmospheric oxygen conditions. Primary ileal intestinal epithelial stem cells (IECs) were expanded and maintained as 3D spheroid cultures in Matrigel (BD Biosciences) and 50% L-WRN conditioned medium (CM) containing 10 μM Y-27632 ROCK inhibitor (Tocris Bioscience), and 10 μM SB431542 (Selleckchem) as described previously (53). L-WRN-CM was quality controlled from batch to batch using recently defined methods (11). The medium was changed every 2 days, and the cells were passaged every 7 days using 5e5 cells for a full 12-wells plate. IEC lines were determined to be mycoplasma-negative using the e-Myco plus kit (iNtRON Biotechnology). For all experiments in this study, IECs were used between passages 20 and 30. Human Intestinal Myofibroblasts (Lonza) were maintained in SmGM 2 Smooth Muscle Cell Growth Medium -2 Bulletkit (Lonza). Cells were passaged every 3 days in a 1:5 split. HCT-8 human colorectal adenocarcinoma cells from a male patient (ATCC) were maintained in RPMI 1640 ATCC Modification medium (Thermo Fisher Scientific) supplemented with 10% fetal bovine serum (Sigma). Cells were passaged twice a week at a 1:5 – 1:20 split.

## METHOD DETAILS

### Generating the air-liquid interface human intestinal epithelial cell culture system

#### Irradiating human intestinal myofibroblasts

Human Intestinal Myofibroblasts (HIMF) (Lonza) were trypsinized, suspended in growth medium, and irradiated at 5,000 rads using the Small Animal Radiation Research Platform (SARRP, Xstrahl) at Washington University School of Medicine, St. Louis. After cell viability was assessed with Trypan Blue staining (Thermo Fisher), irradiated cells were quantified, aliquoted in freezing medium (growth medium with 10% FBS and 10% DMSO) and stored at -80°C for short-term use (weeks) or in liquid nitrogen for long-term use (months).

#### Seeding irradiated HIMF feeder cell layer on transwells

Transwells (polyester membrane, 0.4 mm pore; Corning Costar) were coated with Matrigel (Corning) diluted 1:10 in cold PBS, then incubated at 37°C for 20 min. Excess Matrigel solution was aspirated immediately before adding irradiated HIMF (iHIMF) cells. HIMF cells were thawed, resuspended in growth medium, and seeded onto transwells at 8x10^4^ cells/transwell. Growth medium was added to the top and bottom of the transwell and incubated at 37°C for approximately 24 hr before seeding the hIEC spheroids.

#### Seeding the human intestinal epithelial cell monolayers and creating air-liquid interface

Human ileal spheroids were isolated from biopsies obtained from de-identified healthy donors though the BioBank Core of the Digestive Diseases Research Core Center, Washington University School of Medicine in St. Louis. Human ileal spheroids from 7-day-old stem cell cultures were recovered from Matrigel and dissociated with trypsin as described previously(54). Cells were quantified, suspended in 50% CM with 10 μM Y-27632 and 10 μM SB431542, and plated onto iHIMF monolayers at 5x10^4^ cells/transwell. HIMF growth medium was replaced with 50% CM with 10 μM Y-27632 and 10 μM SB431542 in both the top and bottom compartments for two days, then replaced with 50% CM with 10 μM Y-27632 only and replenished every other day. After 7 days of culture, the medium in the top compartment was removed to create the air-liquid interface (ALI). For *Cryptosporidium* growth experiments, 10 μM of SB431542 was added back to bottom chamber medium on the day of top medium removal and for the rest of the experiment. Medium in the bottom compartment of the transwell continued to be changed every other day. Liquid and/or mucus that appeared in top compartment was removed by aspiration every other day.

### *Cryptosporidium* oocyst preparation and infection of ALI transwells

Before infection, purified *Cryptosporidium* oocysts were treated with a 40% bleach solution (commercial laundry bleach containing 8.25% sodium hypochlorite) diluted in Dulbecco’s Phosphate Buffered Saline (DPBS; Corning Cellgro) for 10 min on ice. Oocysts were then washed 4 times in DPBS containing 1% (wt/vol) bovine serum albumin (BSA) (Sigma) before resuspending with 1% BSA. Oocysts were added to monolayers in 30 μl of 50% CM three days post top medium removal. After 4 hr, monolayers were washed twice with DPBS to remove extracellular parasites and re-establish the air-liquid interface.

### Preparation, staining and imaging of flat mounted transwell membranes

Transwells were moved to a new 24-well plate with PBS in the bottom chamber. Monolayers were fixed by adding 100 μl 4% paraformaldehyde (Polysciences) for 15 min. Cells were washed twice with PBS and then permeabilized and blocked with DPBS containing 1% BSA and 0.1% Triton X-100 (Sigma) for 20 min. Primary antibodies were diluted in blocking buffer for staining: 1B5 and 1A5 (purified mouse mAbs) were used at 1:500, pan Cp (rabbit pAb) was used at 1:1000, Crypt-a-glo TM (mouse mAb, Waterborne, Inc) was used at 1 drop per transwell, and 4D8 (hybridoma supernatant) was used at 1:40. Cells were incubated with primary antibodies for 60 min at room temperature, washed twice with PBS, then incubated for 60 min at room in secondary antibodies conjugated to Alexa Fluor dyes (ThermoFisher Scientific) diluted 1:500 in blocking buffer. Nuclear DNA was stained with Hoechst 33342 (ThermoFisher Scientific) diluted 1:1000 in blocking buffer for 10 mins at room temperature, then the membrane was cut out of the transwell insert using a scalpel and mounted on a glass slide with Prolong Diamond Antifade Mountant (ThermoFisher Scientific). For EdU staining, 10 mM EdU was added to medium in the bottom chamber of the transwell overnight. The transwell was then fixed with 4% paraformaldehyde and permeabilized as described above. EdU was labeled with the Click-iT Plus EdU Alexa Fluor 594 (Thermo Fisher Scientific) Imaging Kits. Primary and secondary antibody staining were done after EdU labeling. Imaging was performed on a Zeiss LSM880 confocal laser scanning microscope (Carl Zeiss Inc.) equipped with a Plan-Apochromat 63X (NA 1.4) DIC objective and a photo multiplier tube (PMT) detector. Images were acquired using ZEN black software (version 2.1 SP3) and manipulated in FIJI(55) and GIMP(56).

### Preparation, staining and imaging of transversely sectioned transwell membranes

Transwells were treated with 4% paraformaldehyde in both top and bottom chambers for 20 min at room temperature, washed three times in 70% ethanol, then incubated for 20 min in 70% ethanol (top and bottom chambers). The transwell membranes were cut from the insert using a scalpel and embedded in 1% agar and then processed for paraffin embedding. For hematoxylin and eosin (H&E) staining and immunohistochemistry, 5 mm transverse sections were cut and processed for staining following standard procedures of the Digestive Disease Research Core Center (DDRC, Washington University in St. Louis). Sections were imaged using a Zeiss Observer.D1 inverted wide-field fluorescence microscope with Axiocam 503 dual B/W and color camera or a Zeiss Axioskop Mot Plus fluorescence microscope equipped with a 100X, 1.4 N.A. Zeiss Plan Apochromat oil objective and an AxioCam MRm monochrome digital camera. For immunostaining, slides were deparaffinized, and antigen retrieval was performed with Trilogy (Sigma-Aldrich). Slides were then stained as described above using the following antibodies: mouse anti-β-catenin (1:500, BD Bioscience), and mouse anti-E-cadherin (1:500, BD Bioscience) with goat anti-mouse IgG Alexa Fluor 488 (Thermo Fisher Scientific); mouse anti-mucin 2 (1:100, SantaCruz) with goat anti-mouse IgG Alexa 568; rabbit anti-ZO-1 (1:500, Invitrogen) and rabbit anti-lysozyme (1:500, Novus) with goat anti-rabbit IgG Alexa Fluor 568 (Thermo Fisher Scientific); rabbit anti-villin (1:1000, Abcam), rabbit anti-Chromogranin A antibody (1:100, Abcam), and rabbit anti-ZO-1 (1:500, Invitrogen) with goat anti-rabbit IgG Alexa Fluor 488 (Thermo Fisher Scientific); and Hoechst 33342 (Invitrogen) for nuclear staining. Images were taken using a 20x (N.A. 1.30) or or a 63X oil objective on a Zeiss Axioskop 2 equipped for epifluorescence. Images were acquired using Axiovision software (Carl Zeiss Inc.) and manipulated in FIJI (55) and GIMP (56).

### Measuring *Cryptosporidium* growth and host cell viability by qPCR

To monitor infection by quantitative PCR, DNA was collected and purified from infected transwells using the QIAamp DNA Mini kit (Qiagen). Briefly, 100 μl Trypsin-EDTA solution (Sigma-Aldrich) was added to monolayer and incubated at 37°C for 5 min, before quenching of trypsin with 100 μl 50% CM. Then, cells were scraped using a pipette tip and stored at -20°C in 1.7 ml tubes until further processing. Samples were centrifuged at 3,000 rpm for 3 min, then the supernatant was discarded.

Cells were resuspended and incubated in Buffer ATL and proteinase K (both reagents provided by kit) in a 56°C water bath for 3-24 hr before column purification. Purified DNA was eluted in 100 μl Buffer AE then diluted 1:5 with H_2_O. Two μl of the diluted DNA was used as template in a qPCR reaction with SYBR Green JumpStart Taq ReadyMix (Sigma-Aldrich). Primer sequences targeting *Cryptosporidium* GAPDH were as follows: forward primer 5’-CGGATGGCCATACCTGTGAG-3’ and reverse primer 5’- GAAGATGCGCTGGGAACAAC-3’. A standard curve for *Cryptosporidium* genomic DNA was generated by purifying DNA from a known number of oocysts and creating a dilution series. Primer sequences targeting human GAPDH were as follows: forward primer 5’- TGAGTACGTCGTGGAGTCCA-3’ and reverse primer 5’-CTGAGTCAGCTTCCCCTCC-3’. A standard curve for human genomic DNA was generated by purifying DNA from a known number of human ileal stem cells and creating a dilution series. Reactions were performed on a QuantStudio 3 Real-Time PCR System (ThermoFisher) with the following amplification conditions: priming at 95°C for 2 min then 40 cycles of denaturing at 95°C for 10 sec, annealing at 60°C for 20 sec, extension at 72°C for 30 sec, followed by a melt curve analysis to detect non-specific amplification. Genomic DNA equivalents in each sample were determined by the QuantStudio Design & Analysis software based on the standard curve samples present on each plate.

### Measuring trans-epithelial electrical resistance across hALI and HCT-8 cultures, FITC-dextran permeability assay and intercellular junction organization quantification measurement

hALI transwells were infected with *Cryptosporidium* oocysts or mock-infected as described above, 3 days after top media removal. HCT-8 cells were seeded on transwells coated with Matrigel and seeded with HIMF cells 24 hr before, then were incubated for 3 days in in RPMI 1640 ATCC Modification medium (Thermo Fisher Scientific A1409101) supplemented with 10% fetal bovine serum (Sigma) (HCT-8 medium). Cells were then infected by adding 100 μl of suspension of 10^5^ bleached oocysts in HCT-8 medium, or mock-infected by adding 100 μl of HCT-8 medium in the top chamber.

After 4 hr, top-chamber medium was removed, the monolayer washed 3 times, then 100 μl of HCT-8 medium was added back. Electrical resistance was measured 4 hr and 24 hr after infection, with a calibrated Millicell-ERS voltohmmeter (Millipore) at room temperature. For dextran flux experiments, 2 mg/ml 3-5kDa fluorescein dextran (Sigma) in 50% CM with 10 μM Y-27632, or, in HCT-8 medium was placed in the upper transwell and incubated for 20 min at 37°C, then medium was harvested from the bottom chamber. Concentrations of FITC-dextran in samples were measured using a Cytation 3 Image Reader (BioTek), with technical duplicates for each sample, and compared to a standard curve generated from dilution standards. Disruption of adherens junctions and tight junctions was quantified using intercellular junction organization quantification (IJOQ), a newly described(57) automated algorithm which analyzes and quantifies the degree of organizational disruption of intercellular junctions. To do so, transwells were harvested 24 hr after *C. parvum* infection or mock-infection. HCT- 8 cells were grown on transwells without Matrigel of HIMFs (E-cadherin), or on non-coated coverslips (ZO-1) for 3 days in HCT-8 medium before infection, then harvested 24 hr after infection. Cells were fixed, stained for ZO-1 or E-cadherin, mounted, and images captured by confocal microscopy as described above. Transwells harvested from non-infected cells were used for calibration of the program. Six different fields of the same sample were captured and submitted to the program to obtain a mean value based on these six images for each sample.

### RNA sequencing and secreted protein quantification of infected vs. non-infected ALI

#### RNA and medium collection and RNA purification

Cells were infected 10 days after seeding, i.e. 3 days after top medium removal, as described above. RNA was harvested from *C. parvum*-infected, *C. hominis*-infected or mock-infected transwells 24 hr after infection. To collect RNA, transwells were incubated in 100 μl trypsin-EDTA solution at 37°C for 5 min before quenching of trypsin with 100 μl 50% CM. Samples were then centrifuged at 3,000 rpm for 3 min, the supernatant discarded and cells resuspended in RNAlater Stabilization and Storage solution (ThermoFisher), then stored at -80°C until further processing. Samples were then centrifuges at 3,000 rpm for 3 min, then supernatant discarded, and RNA extracted using the RNeasy Mini Kit (Qiagen) according to manufacturers guidelines. Two transwells were combined per column for each condition, from three independent experiments for a total of 9 samples. Concurrently, medium from the bottom chamber was harvested 4, 24 and 72 hr after infection, and stored at -80°C until further processing for quantification of soluble secreted chemokines. Two transwells were combined at each timepoint from three independent experiments for a total of 27 samples.

#### Library preparation, RNA sequencing and RNA-Seq data analysis

Total RNA integrity was determined using Agilent Bioanalyzer or 4200 Tapestation. Library preparation was performed with 500ng to 1 μg of total RNA. Ribosomal RNA was removed by an RNase-H method using RiboErase kits (Kapa Biosystems). mRNA was then fragmented in reverse transcriptase buffer and heating to 94°C for 8 min. mRNA was reverse transcribed to yield cDNA using SuperScript III RT enzyme (Life Technologies, per manufacturer’s instructions) and random hexamers. A second strand reaction was performed to yield ds-cDNA. cDNA was blunt ended, had an A base added to the 3’ ends, and then had Illumina sequencing adapters ligated to the ends. Ligated fragments were then amplified for 12-15 cycles using primers incorporating unique dual index tags. Fragments were sequenced on an Illumina NovaSeq-6000 using paired end reads extending 150 bases. Mean sequencing depth was approximately 29.6 million reads per sample (range of 19.7-34.7 million). Raw reads were mapped to the human genome (GRCh38 assembly, Ensembl) using Kallisto, version 0.46. The quality of raw reads, as well as results of Kallisto mapping, were summarized using fastqc and multiqc. Mean percentage of reads mapping was 73.5% (range of 68.2%-78.0%). Transcript-level expression data were imported into the R environment (version 4.0.4; R Foundation for Statistical Computing) and summarized to the gene-level using TxImport with annotations provided by BioMart.

Filtering was carried out to remove lowly expressed genes, and data were normalized using the Trimmed Mean of M values method in EdgeR. For differential expression analysis, precision weights were applied to each gene based on its mean-variance relationship using the VOOM function from the Limma package. Linear modeling and Bayesian statistics were employed via Limma to identify genes that were up or down regulated in the infected transwells, compared to not infected, by 2-fold or more, with a Benjamini–Hochberg adjusted *P* value (false-discovery rate) of less than or equal to .05.

#### Medium samples preparation and quantification of soluble secreted chemokines

Samples were thawed on ice, centrifuged at 15,000 rcf for 5 min at 4°C to remove particulates, then 50 μl of sample or standard was added per well (in duplicate) with premixed beads in a 96-well plate according to the manufacturer instructions (ThermoFisher Procartaplex Human Simplex kit). The plate was incubated overnight on a shaker at 4°C, and the final detection steps and machine analysis were performed the next morning. The beads were read for MFI of streptavidin-PE using a FLEXMAP3D Luminex (Luminex Corp) machine. The analysis software MilliporeSigma Belysa v.1.0 was used to calculate the pg/ml for each analyte from the standard curve using a 5-parameter logistical fit algorithm.

### CSpV1 quantification in oocysts

Three samples of the two *Cryptosporidium* strains used in the present study, produced in, and purified from 3 different animals (calves for *C. parvum*, piglet for *C. hominis*) on different days, using similar methods, were analyzed for CSpV1 load. Oocysts were bleached as described above, then submitted to excystation by being incubated for 1 hr at 37°C suspended in 1.5% taurocholic acid. Sporozoites were centrifuged 3 min at 2500 rpm, the supernatant discarded, and sporozoites washed twice, before proceeding to RNA extraction and cDNA synthesis as described above. CSpV1 virus was quantified by qPCR and normalized on CpGAPDH, using the following primers targeting the capsid protein gene: CSpV1 Forward: 5’-TGCAGTTTACTATCCAGTGG-3’; CSpV1 Reverse 5’-GCAGAAGGGTTCTATGATTC-3’. CpGAPDH primers are described above.

### Rotavirus preparation, infection and quantification

African Green Monkey kidney epithelial cell line MA104 (CRL-2378.1) was obtained from American Type Culture Collection (ATCC) and cultured in Medium 199 (Gibco) supplemented with 10% heat-inactivated fetal bovine serum (VWR), 100 units/ml penicillin and 100 μg/ml streptomycin. Rotavirus strains were propagated as describes previously (58) and the titers of virus concentrate was determined in MA104 cells using a standard plaque assay (59). Rotavirus inoculum was prepared by activating the stock with 5 μg/ml trypsin for 20 min at 37°C and diluting it into 50% CM with 10 μM Y- 27632 up to the total volume of 100 μl, with 50 μg/ml soybean trypsin inhibitor. The inoculum was carefully added to the upper compartment of IEC transwell culture and was incubated at 37°C. After 2 hr, the inoculum was removed by aspiration. Cells were harvested and RNA extracted as described above, then retrotranscripted into cDNA using the iScript cDNA Synthesis Kit (Bio-Rad) according to manufacturer recommendations. To quantify RV NSP5 and cellular IFNL3 transcripts, we used 2x TaqMan Fast Advanced master mix (Applied Biosystems) and 2x SYBR Green master mix (Applied Biosystems) in 25 μl reaction volume, respectively. Both transcript levels were normalized to the level of housing keeping gene GAPDH, which was also measured using 2x SYBR Green master mix. The sequences of primers used are as follows: GAPDH Forward: 5’-GGAGCGAGATCCCTCCAAAAT-3’, Reverse: 5’-GGCTGTTGTCATACTTCTCATGG-3’; Rotavirus NSP5 Forward: 5’- CTGCTTCAAACGATCCACTCAC-3’, Reverse: 5’-TGAATCCATAGACACGCC-3’, Probe: 5’-Cy5 dye- TCAAATGCAGTTAAGACAAATGCAGACGCT-Iowa Black RQ quencher-3’; IFNL3 Forward: 5’- TAAGAGGGCCAAAGATGCCTT-3’, Reverse: 5’-CTGGTCCAAGACATCCCCC-3’

### Quantification and statistical analysis

All statistical analyses were performed in the R environment using the rstatix package (version 0.7.0). When comparing the means of two groups, we used the Student’s t-test. When comparing the means of two or more groups across time, we used a one-way ANOVA followed by multiple comparisons using the Tukey’s HSD test. Non-parametric tests were used when data did not pass a Shapiro-Wilk test for normality or sample sizes were low. A Mann-Whitney U test was used when comparing the mean of two groups, while a Kruskal-Wallis test followed by a Dunn’s multiple comparisons test was used when comparing the means of one variable across three or more groups. Statistical parameters for each experiment including statistical test used, technical replicates (n) and independent biological replicates (N) are reported in the figure legends.

### Blinding and inclusion/exclusion

Samples were not blinded for analysis although they were repeated independently as stated in the text and figure legends. All samples were included in the analysis.

## DATA AND SOFTWARE AVAILABILITY

Raw RNA-seq reads and analyzed data generated in this study have been deposited in the Gene Expression Omnibus database under accession numbers GSE241570 (*C. hominis*) & GSE241571 (*C. parvum*).

## AUTHOR CONTRIBUTIONS

**Conceptualization:** Valentin Greigert, Siyuan Ding, L. David Sibley. **Data curation:** Valentin Greigert, Iti Saraav, Juhee Son. **Formal Analysis:** Valentin Greigert, Iti Saraav. **Investigation:** Valentin Greigert, Iti Saraav, Juhee Son. **Visualization:** Valentin Greigert. **Writing – original draft:** Valentin Greigert. **Resources:** Denise Dayao, Avan Antia, Saul Tzipori, William H. Witola, Thaddeus S. Stappenbeck. **Supervision:** Siyuan Ding, Saul Tzipori, L. David Sibley. **Writing – review & editing:** Valentin Greigert, Siyuan Ding, L. David Sibley.

## ACKNOWLEDGMENTS

We thank Soumya Ravindran for initial studies and Kelli VanDussen and Yi Wang for helpful advice on developing the culture methods used here. Select illustrations were kindly provided by Abigail Kimball in association with InPrint at Washington University in St. Louis. Sequencing was performed by the Genome Technology Access Core, McDonnell Genome Institute, Washington University. Imaging was performed using the Molecular Microbiology Imaging Facility, Washington University School of Medicine. Histological services and spheroid cell lines were provided by the Digestive Diseases Research Core Center (NIDDK P30 DK052574), Washington University School of Medicine. Rotarix (RV1) and Rotateq (RV5) stocks were a kind gift from Kristen Ogden at Vanderbilt University.

Immunology assays were conducted with support of the Bursky Center for Human Immunology and Immunotherapy Programs at Washington University, Immuno-monitoring Laboratory. Support for studies in the Sibley laboratory was provided by a grant from Open Philanthropy. Support for studies conducted in the Ding lab were provided by NIH T32 DK007130, and R01 AI150796.

---

**Fig. S1.**
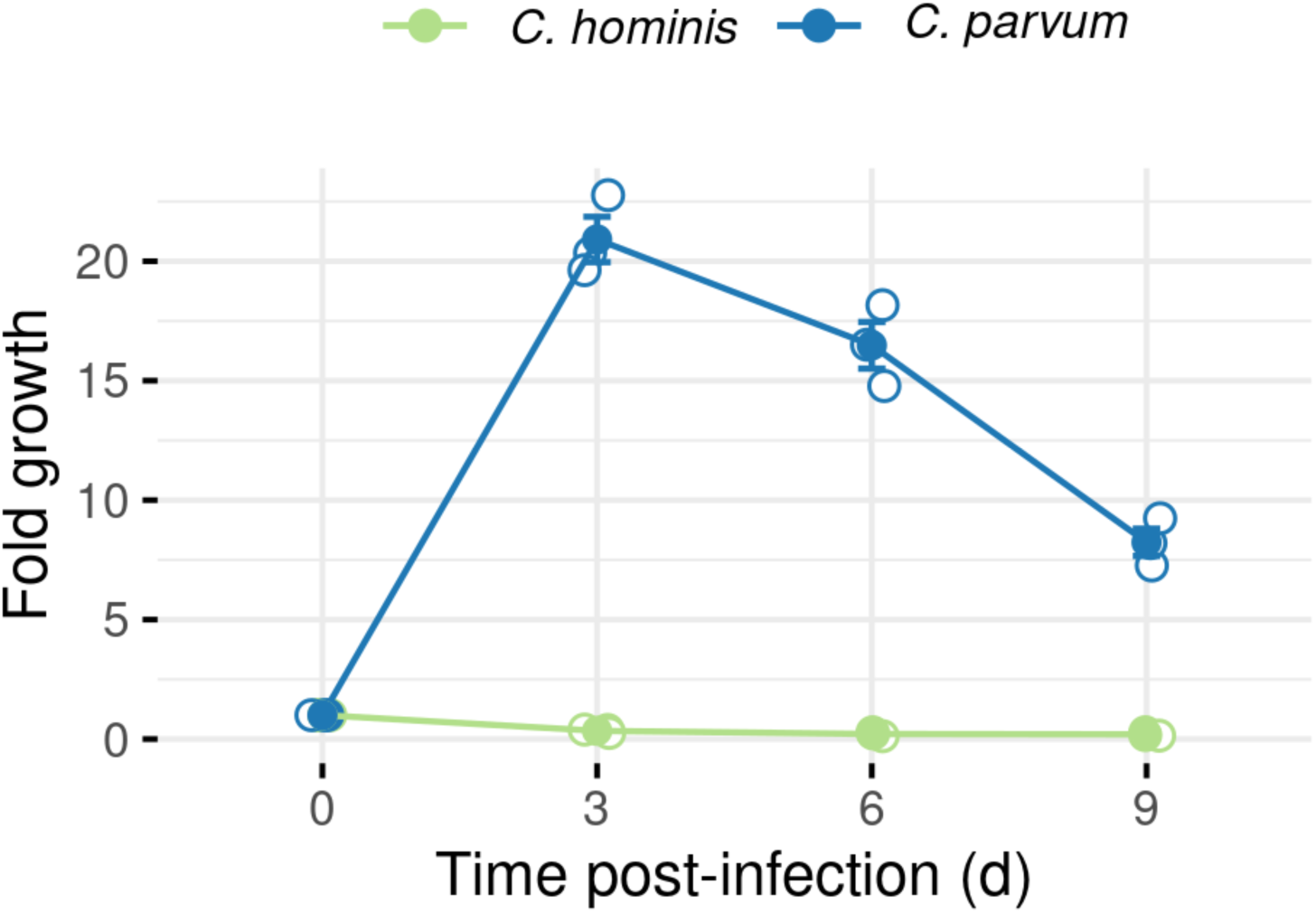
Growth of *C. parvum* and *C. hominis* in mouse ALI cultures over a 9-day period, as measured by qPCR. Mouse ALI does not sustain *C. hominis* growth. Mean ± S.E. N = 1, n = 3.

**Fig. S2.**
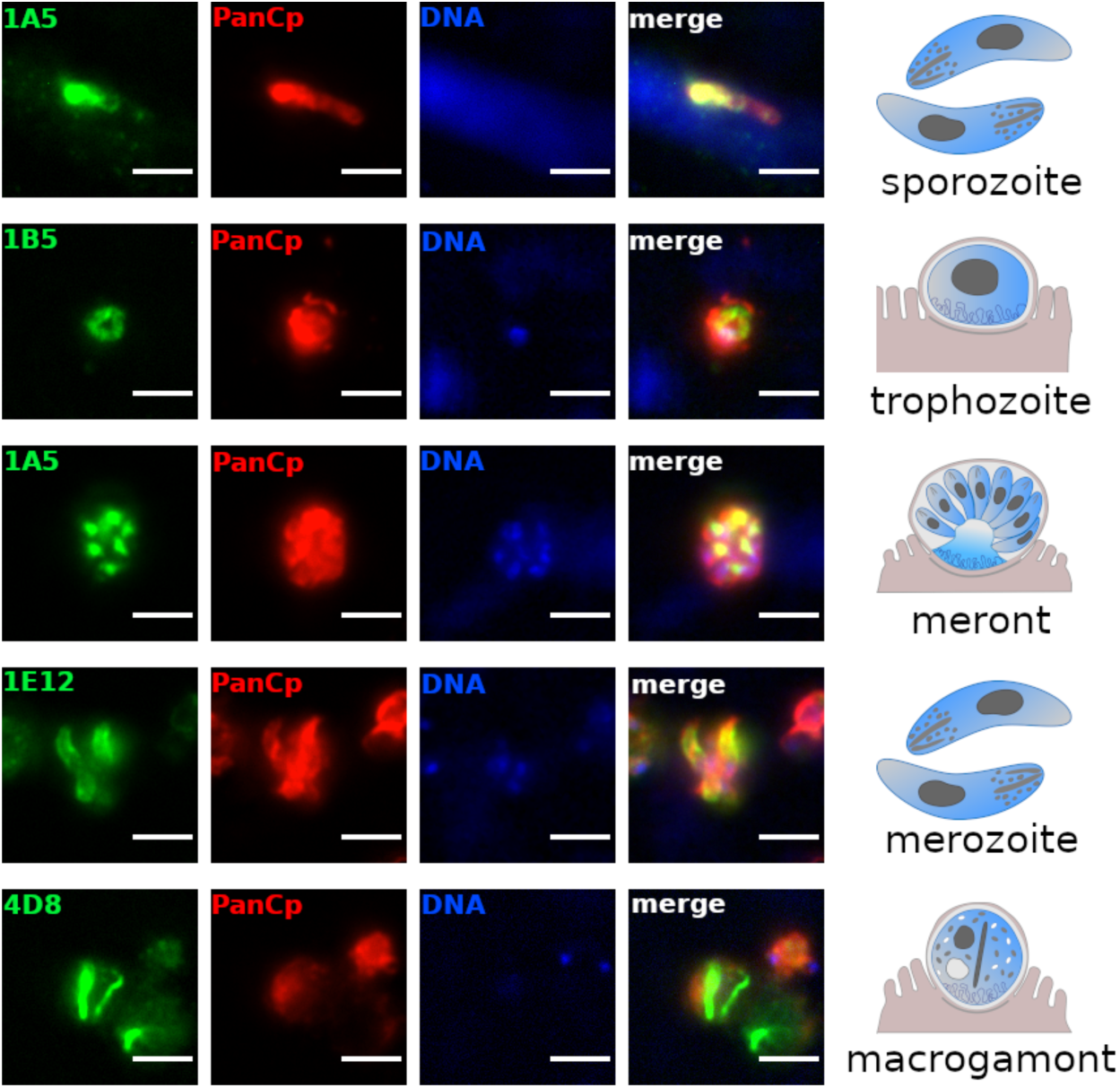
Immunostaining of various stages of *C. hominis* cultured on HCT-8 cells using a panel of previously described mAb to *C. parvum* (green) and a pan-*Cryptosporidium* polyclonal antibody (red). Scale bar = 3 μm.

**Fig. S3.**
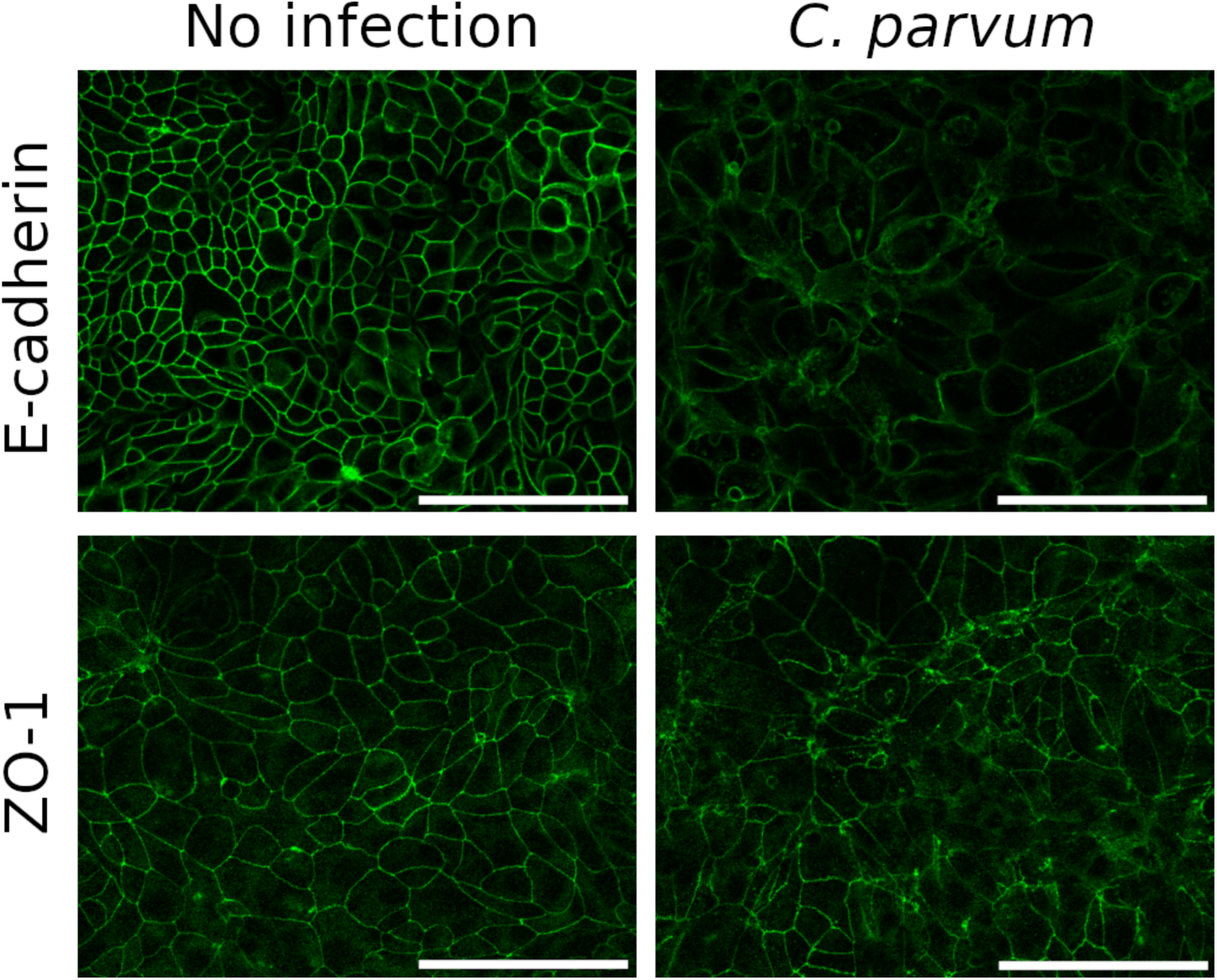
Immunostaining of HCT-8 transwell (E-cadherin) or coverslips (ZO-1) cultures showing tight junction (TJ; ZO-1, green) and adherens junction (AJ; E-cadherin, green) organization, with or without infection with *C. parvum* (not stained). Scale bar = 50 μm.

**Fig. S4.**
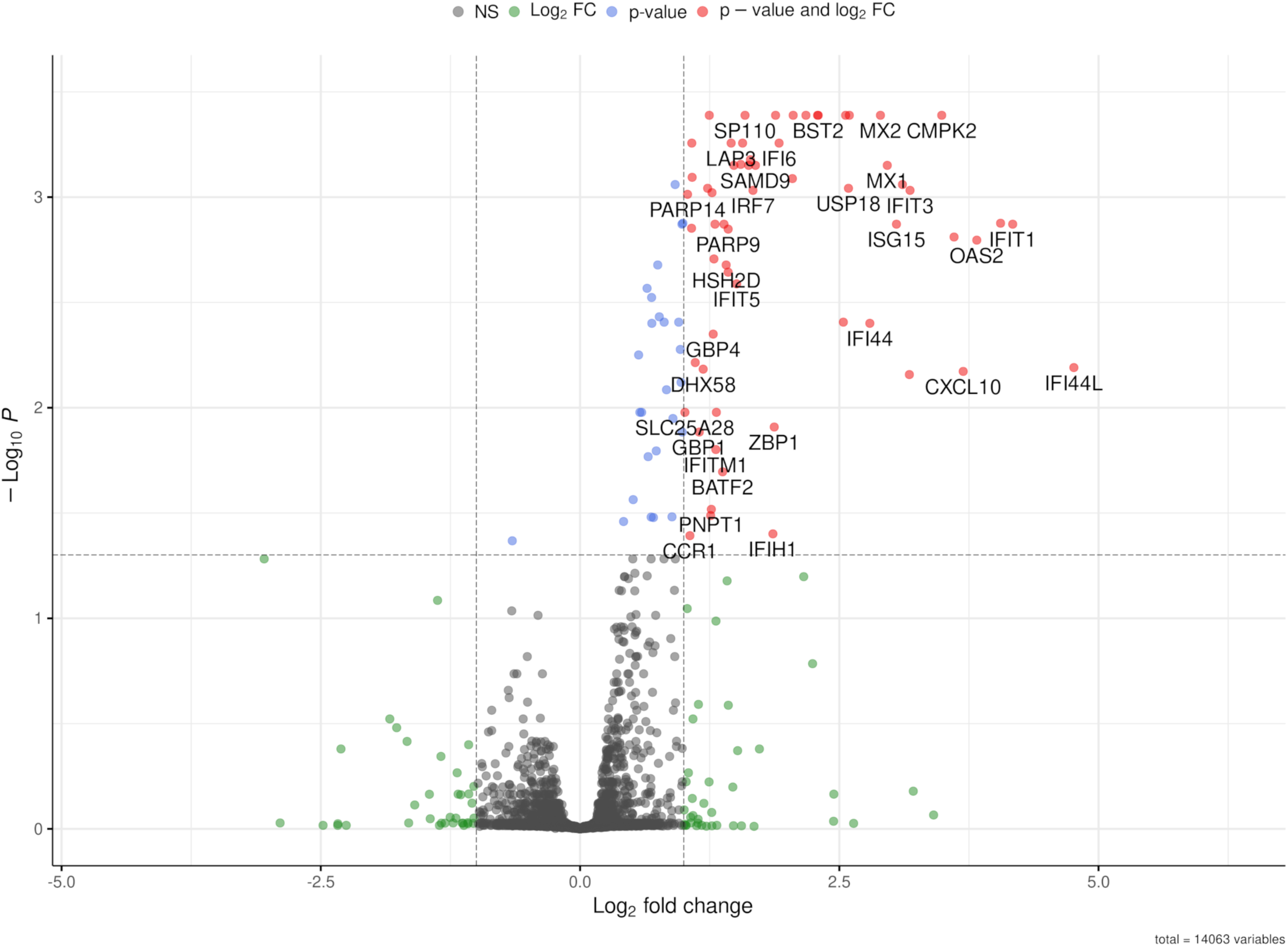
Volcano plot of differentially expressed genes in hALI 24 hr after infection with *C. hominis*. Genes up- or down-regulated by more than 2-fold and *P* < 0.05 are shown in red.

## REFERENCES

1. Feng Y, Ryan UM, Xiao L. Genetic Diversity and Population Structure of *Cryptosporidium*. Trends Parasitol. 2018;34(11):997–1011.

2. Kotloff KL, et al. Burden and aetiology of diarrhoeal disease in infants and young children in developing countries (the Global Enteric Multicenter Study, GEMS): a prospective, case-control study. The Lancet. 2013;382(9888):209–222.

3. Kotloff KL. The Burden and Etiology of Diarrheal Illness in Developing Countries. Pediatr Clin North Am. 2017;64(4):799–814.

4. Checkley W, et al. A review of the global burden, novel diagnostics, therapeutics, and vaccine targets for *Cryptosporidium*. The Lancet Infectious Diseases. 2015;15(1):85–94.

5. Theodos CM, et al. Profiles of healing and nonhealing *Cryptosporidium parvum* infection in C57BL/6 mice with functional B and T lymphocytes: the extent of gamma interferon modulation determines the outcome of infection. Infect Immun. 1997;65(11):4761–4769.

6. Rehg JE. Effect of interferon-gamma in experimental *Cryptosporidium parvum* infection. J Infect Dis. 1996;174(1):229–232.

7. Lee S, Beamer G, Tzipori S. The piglet acute diarrhea model for evaluating efficacy of treatment and control of cryptosporidiosis. Hum Vaccin Immunother. 2019;15(6):1445–1452.

8. Dayao DA, et al. An immunocompetent rat model of infection with *Cryptosporidium hominis* and Cryptosporidium parvum. Int J Parasitol. 2020;50(1):19–22.

9. Bhalchandra S, Lamisere H, Ward H. Intestinal organoid/enteroid-based models for *Cryptosporidium*. Current Opinion in Microbiology. 2020;58:124–129.

10. VanDussen KL, Sonnek NM, Stappenbeck TS. L-WRN conditioned medium for gastrointestinal epithelial stem cell culture shows replicable batch-to-batch activity levels across multiple research teams. Stem Cell Research. 2019;37:101430.

11. Wilke G, et al. A Stem-Cell-Derived Platform Enables Complete *Cryptosporidium* Development In Vitro and Genetic Tractability. Cell Host Microbe. 2019;26(1):123–134.e8.

12. Wilke G, et al. Monoclonal Antibodies to Intracellular Stages of *Cryptosporidium parvum* Define Life Cycle Progression In Vitro. mSphere. 2018;3(3). 10.1128/mSphere.00124-18

13. Griffiths JK, et al. *Cryptosporidium parvum* infection of Caco-2 cell monolayers induces an apical monolayer defect, selectively increases transmonolayer permeability, and causes epithelial cell death. Infect Immun. 1994;62(10):4506–4514.

14. de Sablet T, et al. *Cryptosporidium parvum* increases intestinal permeability through interaction with epithelial cells and IL-1β and TNFα released by inflammatory monocytes. Cell Microbiol. 2016;18(12):1871–1880.

15. Kumar A, et al. *Cryptosporidium parvum* disrupts intestinal epithelial barrier function via altering expression of key tight junction and adherens junction proteins. Cell Microbiol. 2018;20(6):e12830.

16. Priyamvada S, et al. *Cryptosporidium parvum* infection induces autophagy in intestinal epithelial cells. Cell Microbiol. 2021;23(4):e13298.

17. Sateriale A, et al. The intestinal parasite *Cryptosporidium* is controlled by an enterocyte intrinsic inflammasome that depends on NLRP6. Proc Natl Acad Sci U S A. 2021;118(2). 10.1073/pnas.2007807118

18. Leoni F, et al. Characterisation of small double stranded RNA molecule in *Cryptosporidium hominis*, *Cryptosporidium felis* and *Cryptosporidium meleagridis*. Parasitol Int. 2006;55(4):299–306.

19. Nibert ML, et al. Cryspovirus: a new genus of protozoan viruses in the family Partitiviridae. Arch Virol. 2009;154(12):1959–1965.

20. Ding S, et al. Rotavirus VP3 targets MAVS for degradation to inhibit type III interferon expression in intestinal epithelial cells. Elife. 2018;7:e39494.

21. Cardenas D, et al. Two- and Three-Dimensional Bioengineered Human Intestinal Tissue Models for *Cryptosporidium*. Methods Mol Biol. 2020;2052:373–402.

22. Heo I, et al. Modelling *Cryptosporidium* infection in human small intestinal and lung organoids. Nat Microbiol. 2018;3(7):814–823.

23. Lamisere H, et al. Differential Response to the Course of *Cryptosporidium* parvum Infection and Its Impact on Epithelial Integrity in Differentiated versus Undifferentiated Human Intestinal Enteroids. Infect Immun. 2022;90(11):e0039722.

24. Ferguson SH, et al. Interferon-λ3 Promotes Epithelial Defense and Barrier Function Against *Cryptosporidium parvum* Infection. Cell Mol Gastroenterol Hepatol. 2019;8(1):1–20.

25. Tang X, et al. Specific and Non-Specific Adhesion in Cancer Cells with Various Metastatic Potentials. In: Wagoner Johnson A, Harley BAC, eds. Mechanobiology of Cell-Cell and Cell-Matrix Interactions. Boston, MA: Springer US; 2011:105–122

26. Crawford CK, Kol A. The Mucosal Innate Immune Response to *Cryptosporidium parvum*, a Global One Health Issue. Front Cell Infect Microbiol. 2021;11:689401.

27. Chen W, et al. Gamma interferon functions in resistance to *Cryptosporidium parvum* infection in severe combined immunodeficient mice. Infect Immun. 1993;61(8):3548–3551.

28. Mead JR, You X. Susceptibility differences to *Cryptosporidium parvum* infection in two strains of gamma interferon knockout mice. J Parasitol. 1998;84(5):1045–1048.

29. Hayward AR, Chmura K, Cosyns M. Interferon-gamma is required for innate immunity to *Cryptosporidium parvum* in mice. J Infect Dis. 2000;182(3):1001–1004.

30. Leav BA, et al. An early intestinal mucosal source of gamma interferon is associated with resistance to and control of *Cryptosporidium parvum* infection in mice. Infect Immun. 2005;73(12):8425–8428.

31. Gullicksrud JA, et al. Enterocyte–innate lymphoid cell crosstalk drives early IFN-γ-mediated control of *Cryptosporidium*. Mucosal Immunol. 2021;1–11.

32. Pollok RC, et al. Interferon gamma induces enterocyte resistance against infection by the intracellular pathogen *Cryptosporidium parvum*. Gastroenterology. 2001;120(1):99–107.

33. Choudhry N, et al. Dysregulation of interferon-gamma-mediated signalling pathway in intestinal epithelial cells by *Cryptosporidium parvum* infection. Cell Microbiol. 2009;11(9):1354–1364.

34. Gomez Morales MA, et al. Crude extract and recombinant protein of *Cryptosporidium parvum* oocysts induce proliferation of human peripheral blood mononuclear cells in vitro. J Infect Dis. 1995;172(1):211–216.

35. Gomez Morales MA, et al. Cytokine profile induced by *Cryptosporidium* antigen in peripheral blood mononuclear cells from immunocompetent and immunosuppressed persons with cryptosporidiosis. J Infect Dis. 1999;179(4):967–973.

36. White AC, et al. Interferon-gamma expression in jejunal biopsies in experimental human cryptosporidiosis correlates with prior sensitization and control of oocyst excretion. J Infect Dis. 2000;181(2):701–709.

37. Choudhry N, et al. A protective role for interleukin 18 in interferon γ-mediated innate immunity to *Cryptosporidium parvum* that is independent of natural killer cells. J Infect Dis. 2012;206(1):117–124.

38. Ghimire L, et al. The NLRP6 inflammasome in health and disease. Mucosal Immunol. 2020;13(3):388–398.

39. Barakat FM, et al. *Cryptosporidium parvum* infection rapidly induces a protective innate immune response involving type I interferon. J Infect Dis. 2009;200(10):1548–1555.

40. Ye L, Schnepf D, Staeheli P. Interferon-λ orchestrates innate and adaptive mucosal immune responses. Nat Rev Immunol. [published online ahead of print: June 14, 2019]; 10.1038/s41577-019-0182-z

41. Gibson AR, et al. A genetic screen identifies a protective type III interferon response to *Cryptosporidium* that requires TLR3 dependent recognition. PLoS Pathog. 2022;18(5):e1010003.

42. Fitzgerald KA, et al. IKKε and TBK1 are essential components of the IRF3 signaling pathway. Nat Immunol. 2003;4(5):491–496.

43. Deng S, et al. *Cryptosporidium* uses CSpV1 to activate host type I interferon and attenuate antiparasitic defenses. Nat Commun. 2023;14(1):1456.

44. Villares M, Berthelet J, Weitzman JB. The clever strategies used by intracellular parasites to hijack host gene expression. Semin Immunopathol. 2020;42(2):215–226.

45. Zhang X, et al. Interleukin-22 regulates the homeostasis of the intestinal epithelium during inflammation. International Journal of Molecular Medicine. 2019;43(4):1657–1668.

46. Južnić L, et al. SETDB1 is required for intestinal epithelial differentiation and the prevention of intestinal inflammation. Gut. 2021;70(3):485–498.

47. Crawford SE, et al. Rotavirus infection. Nat Rev Dis Primers. 2017;3(1):1–16.

48. Varghese T, Kang G, Steele AD. Understanding Rotavirus Vaccine Efficacy and Effectiveness in Countries with High Child Mortality. Vaccines. 2022;10(3):346.

49. Carvalho MF, Gill D. Rotavirus vaccine efficacy: current status and areas for improvement. Hum Vaccin Immunother. 2019;15(6):1237–1250.

50. Kuhlenschmidt TB, et al. Inhibition of Calcium-Dependent Protein Kinase 1 (CDPK1) In Vitro by Pyrazolopyrimidine Derivatives Does Not Correlate with Sensitivity of *Cryptosporidium parvum* Growth in Cell Culture. Antimicrob Agents Chemother. 2015;60(1):570–579.

51. Akiyoshi DE, et al. Molecular characterization of ribonucleotide reductase from *Cryptosporidium parvum*. DNA Seq. 2002;13(3):167–172.

52. Chappell CL, et al. *Cryptosporidium hominis*: experimental challenge of healthy adults. Am J Trop Med Hyg. 2006;75(5):851–857.

53. Miyoshi H, et al. Wnt5a Potentiates TGF-β Signaling to Promote Colonic Crypt Regeneration After Tissue Injury. Science. 2012;338(6103):108–113.

54. Moon C, et al. Development of a primary mouse intestinal epithelial cell monolayer culture system to evaluate factors that modulate IgA transcytosis. Mucosal Immunology. 2014;7(4):818–828.

55. Schindelin J, et al. Fiji: an open-source platform for biological-image analysis. Nat Methods. 2012;9(7):676–682.

56. The GIMP Development Team. GIMP https://www.gimp.org/. Accessed March 6, 2023

57. Mo D, et al. Dynamic Python-Based Method Provides Quantitative Analysis of Intercellular Junction Organization During *S. pneumoniae* Infection of the Respiratory Epithelium. Frontiers in Cellular and Infection Microbiology. 2022;12.https://www.frontiersin.org/articles/10.3389/fcimb.2022.865528. Accessed March 2, 2023

58. Hou G, et al. Rotavirus NSP1 Contributes to Intestinal Viral Replication, Pathogenesis, and Transmission. mBio. 2021;12(6):e0320821.

59. Shaw RD, Hempson SJ, Mackow ER. Rotavirus diarrhea is caused by nonreplicating viral particles. J Virol. 1995;69(10):5946–5950.

